# SND1 Impairs Calcium Homeostasis in Right Ventricular Failure by Binding and Destabilizing SERCA2a in Cardiomyocytes

**DOI:** 10.1101/2025.04.14.648853

**Authors:** Lizhe Guo, Xingyang Liu, Lu Wang, Yinqi Li, Xiaocheng Zhu, Xu Cheng, Chunyan Ye, Na Chen, Longyan Li, Gang Qin, Yanfeng Wang, Dandan Wang, Jian Qiu, E Wang

**Affiliations:** Department of Anesthesiology, Xiangya Hospital, Central South University, Changsha, China; Hunan Key Laboratory of Molecular Precision Medicine, Xiangya Hospital, Central South University, Changsha, China; National Clinical Research Center for Geriatric Disorders (Xiangya Hospital), Changsha, China

**Keywords:** Right ventricular failure, SND1, SERCA2a, Calcium homeostasis, Pulmonary arterial hypertension

## Abstract

**Background:** Right ventricular failure (RVF) is a major contributor to a poor prognosis in patients with pulmonary arterial hypertension (PAH); however, its underlying molecular mechanisms remain incompletely understood. We identified a significant upregulation of the cardiac fetal gene SND1 in the right ventricular myocardium of RVF rats. This upregulation may be a key component of cardiac fetal reprogramming and play a critical role in RVF progression. However, the precise molecular mechanisms by which SND1 contributes to RVF remain unelucidated.

**Methods:** Rat models of PAH were established by intraperitoneal injection of monocrotaline (MCT). The pulmonary artery pressure and right ventricular function of rats were assessed using transthoracic echocardiography combined with right heart catheterization. Weighted gene co-expression network analysis (WGCNA) of proteomic data identified a significant association between SND1 upregulation and RVF progression. Calcium transient and contractility measurements in single cardiomyocytes were performed to evaluate the effect of SND1 on sarcoplasmic reticulum (SR) function. Liquid chromatography with mass spectrometry/mass spectrometry analysis combined with co-immunoprecipitation (Co-IP) was used to identify sarcoplasmic/endoplasmic reticulum Ca^2+^-ATPase 2A (SERCA2a) as the protein interacting with SND1. The effect of SND1 on the progression of RVF was validated using cardiomyocyte-specific SND1 conditional knockout rats.

**Results:** Proteomic and molecular analyses revealed that the cardiac fetal gene SND1 is reactivated in RVF and may contribute to disease progression. In vitro, SND1 knockdown in neonatal rat ventricular myocytes (NRVMs) enhanced contractility, improved calcium transient amplitude and velocity, and increased cell survival under calcium overload. Mechanistically, SND1 interacts with SERCA2a and promotes its proteasomal degradation, impairing SR calcium reuptake. SND1 knockdown restored SERCA2a stability. In vivo, cardiomyocyte-specific SND1 knockout improved right ventricular function, reduced SERCA2a degradation and apoptosis, and increased survival in RVF rats.

**Conclusion:** Our study revealed that SND1 reactivation (a fetal gene) contributes to the progression of RVF by promoting SERCA2a degradation and impairing calcium handling. Targeted suppression of SND1 enhances cardiomyocyte contractility and survival, highlighting SND1 as a potential therapeutic target for improving right ventricular function in PAH.

**Clinical perspective:** *What Is New?:*

- This study identifies SND1, a reactivated fetal gene, as a key contributor to right ventricular failure (RVF) progression in pulmonary arterial hypertension (PAH).
- SND1 interacts with SERCA2a, promoting its proteasomal degradation, which disrupts sarcoplasmic reticulum calcium reuptake and weakens cardiomyocyte contractility.
- Cardiomyocyte-specific SND1 deletion restores SERCA2a levels, improves calcium handling, enhances right ventricular function, and increases survival in RVF rats.

*What Are the Clinical Implications?:*

- The reactivation of fetal genes such as SND1 is not just a marker but a pathogenic driver in maladaptive right ventricular remodeling.
- Targeting SND1 represents a novel therapeutic approach to improve right ventricular contractility and prevent heart failure progression in patients with PAH.
- These findings may inform the development of gene- or protein-targeted therapies aimed at stabilizing SERCA2a and preserving right heart function in high-risk populations.

## Introduction

Right ventricular failure (RVF) is a common outcome of advanced pulmonary vascular diseases, parenchymal lung diseases, and left ventricular failure.^1^ RVF results in elevated central venous pressure and sodium and water retention, contributing to adverse clinical outcomes.^2–5^ Increased afterload is a key driver of maladaptive pathological changes in the right ventricular myocardium. Although this pathophysiological mechanism is well recognized, our understanding of the molecular processes underlying the transition from normal right ventricular structure and function to maladaptive and ultimately failing phenotypes remains limited. Furthermore, effective therapeutic strategies targeting this process are lacking.

Cardiac fetal reprogramming, marked by the suppression of adult and reactivation of fetal gene profiles in the diseased myocardium, has emerged as a key mechanism in maladaptive remodeling.^6^ In our previous study, we reported a metabolic shift from fatty acid β-oxidation to glycolysis in the failing right ventricle,^7^ a hallmark of metabolic remodeling that suggests that cardiac fetal reprogramming may also be critically involved in the development and progression of RVF in patients with pulmonary arterial hypertension (PAH).

Staphylococcal Nuclease and Tudor Domain containing 1 (SND1), a fetal gene regulated by mTORC1, is highly expressed in embryonic hearts but silenced in adulthood.^8^ SND1 has been well studied in cancers,^9^ but its role in heart failure remains unexplored. This study shows that SND1 is upregulated in right ventricular cardiomyocytes of RVF rats and is associated with cardiac dysfunction. Knockdown of SND1 in neonatal rat ventricular myocytes (NRVMs) improved calcium handling and contractility and reduced cell death under calcium overload. Mechanistically, SND1 interacts with and promotes the degradation of sarcoplasmic/endoplasmic reticulum Ca^2+^-ATPase 2A (SERCA2a), a key protein in intracellular calcium handling,^10^ impairing SR calcium reuptake. In vivo, cardiomyocyte-specific SND1 deletion improved RV function, reduced apoptosis, and increased survival in PAH rats. Our findings identify SND1 as a novel therapeutic target for improving calcium handling and alleviating RVF.

## Methods and materials

The Supplemental Materials available online provide detailed information on reagents, cell culture conditions, treatments, and experimental procedures. Primer sequences used for PCR, RT-qPCR, and gRNA sequencing are listed in Supplementary Tables S2–S4. The corresponding author can provide the datasets, analytical methods, and study materials upon reasonable request.

## Results

### MCT-induced PAH leads to RVF in rats

To establish a stable PAH model, we injected MCT into the peritoneum of the rats (Figure 1A). The increased intraventricular pressure overload led to progressive tricuspid annular dilation and subsequent tricuspid regurgitation. Therefore, tricuspid regurgitation on Doppler imaging served as a key noninvasive indicator of successfully establishing the PAH model (Figure 1B). We assessed the right ventricular (RV) function by measuring the fractional area change (FAC) of the RV in PAH rats on day 28 post-injection. Compared with the control group, rats in the RVF group exhibited an enlarged right ventricular end-systolic area (RVAs) and a markedly reduced FAC (Figures 1C–1F). Compared with FAC, strain analysis is more sensitive in detecting subtle changes in myocardial function and can detect early abnormalities. Assessments of the global strain (GS) and free wall strain (FWS) revealed that RV strain was significantly impaired in RVF rats compared to controls (Figure 1G–1I). Analysis of key parameters derived from the pressure-volume loops demonstrated a significant increase in RV afterload induced by PAH, accompanied by marked RV dilation (Figure 1J-1M). These results indicate that successful induction of PAH leads to significant RV functional impairment and structural remodeling in rats.

**Figure 1.**
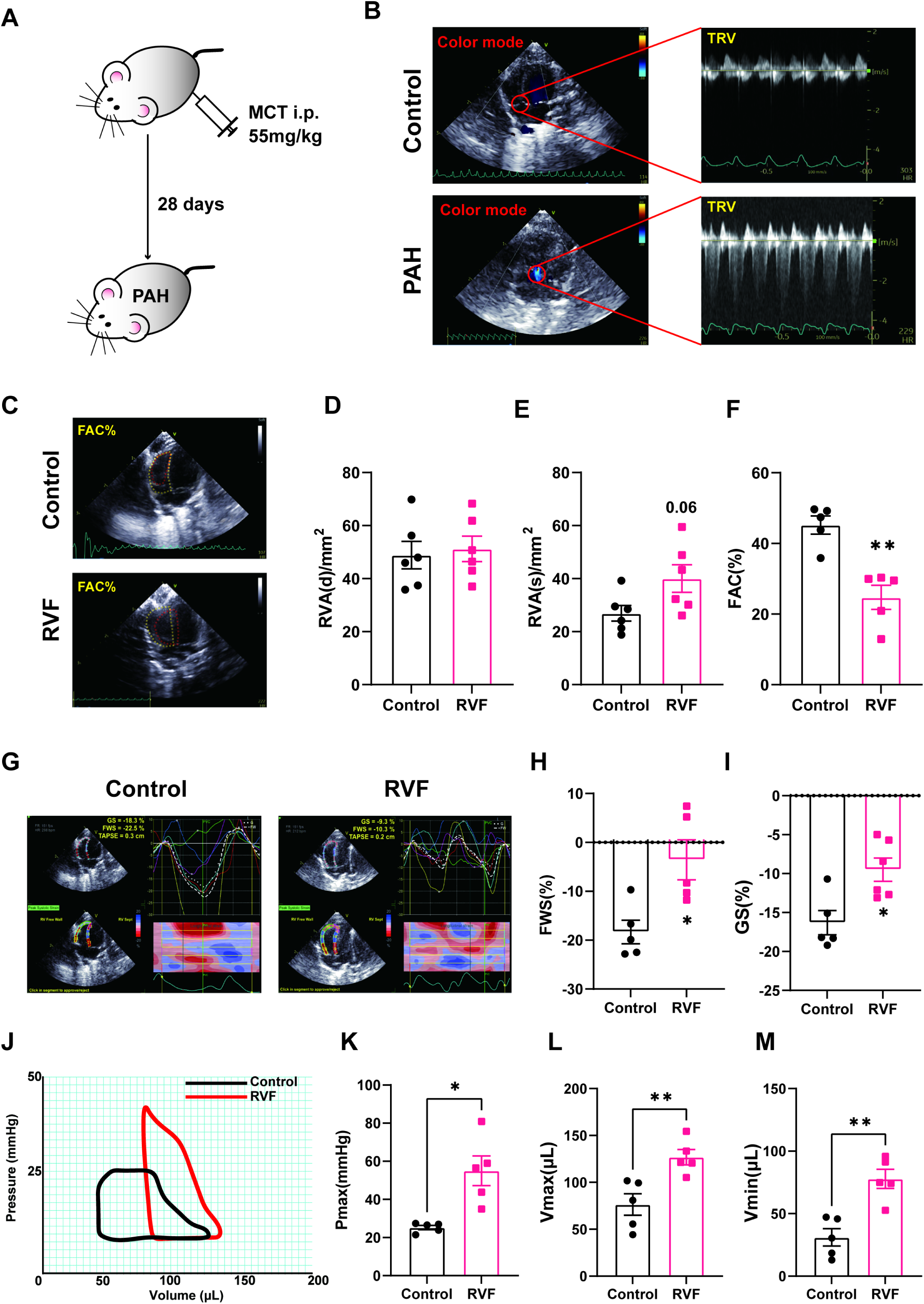
Establishment of the PAH model and confirmation of the RVF phenotype. **A.** Schematic representation of the pulmonary arterial hypertension (PAH) rat model induced by intraperitoneal injection of monocrotaline (MCT). A transthoracic echocardiography was performed to confirm the development of the right ventricular failure (RVF) phenotype. Right ventricular tissues were collected on day 28 for further analysis. **B.** Representative images of color Doppler echocardiography indicating tricuspid regurgitation (TR), which is considered a noninvasive marker of successful PAH model induction. **C–F.** Right ventricular function was assessed by calculating the fractional area change (FAC) using the end-systolic area (RVAs) and the end-diastolic area (RVAd). Results are expressed as the mean ± SEM, n = 6 rats per group. **G.** Representative images of right ventricular strain using speckle-tracking echocardiography. Normal values for ventricular strain are approximately −20, with less negative (i.e., closer to zero) values indicating impaired myocardial contractility. **H.** The absolute value of the free wall strain (FWS) was significantly decreased in RVF rats compared to controls. Results are expressed as the mean ± SEM, n = 6 rats per group. **I.** The absolute value of the global strain (GS) was also significantly reduced in RVF rats. Results are expressed as the mean ± SEM, n = 6 rats per group. **J.** Pressure–volume loop recordings obtained via right heart catheterization were used to assess right ventricular function. **K.** The end-systolic pressure during isovolumetric contraction was significantly elevated in RVF rats compared to controls. Results are expressed as the mean ± SEM, n = 5 rats per group. **L–M.** The right ventricular volume was significantly increased in RVF rats, indicating marked ventricular dilation. Results are expressed as the mean ± SEM, n = 5 rats per group.

### WGCNA links SND1 to RVF progression

In our previous study, proteomic analysis of RV myocardial tissue from RVF rats revealed a metabolic shift toward a fetal-like energy profile, characterized by downregulation of fatty acid β-oxidation and mitochondrial oxidative phosphorylation, along with upregulation of glycolysis-related proteins.^7^ We expanded the sample size for proteomic analysis to further identify key fetal genes involved in the progression of right heart failure. We performed a weighted gene co-expression network analysis (WGCNA) on the dataset.

The sample dendrogram and trait heatmap revealed clear global expression differences between the control and RVF groups and shared intra-group expression characteristics. Among clinical traits, FAC and GS showed the most significant differences between the two groups (Figure 2A). To construct the co-expression network, we selected 14 as the soft threshold to emphasize a strong correlation and make the network conform to the standard scale-free network (Figure 2B). Using the dynamic tree-cut algorithm for hierarchical clustering and subsequent merging of similar expression modules, we identified four distinct co-expression modules: MEblue, MEturquoise, MEgreen, and MEgrey (Figure 2C). Correlation analysis between module eigengenes and clinical traits was visualized in a heatmap (Figure 2D). Notably, the MEblue module was significantly negatively correlated with FAC, suggesting that genes within this module may play important roles in RV systolic dysfunction. Among the MEblue module genes, SND1 exhibited the highest module membership (MM = 0.99355) and a strong correlation with FAC (GS = 0.81552) (Figure 2E). As a newly identified cardiac fetal gene,^8^ SND1 is highly expressed in fetal hearts compared to healthy adult hearts and is markedly upregulated during RVF progression. Our study is the first to identify SND1 as a novel component of cardiac fetal reprogramming in the failing myocardium.

**Figure 2.**
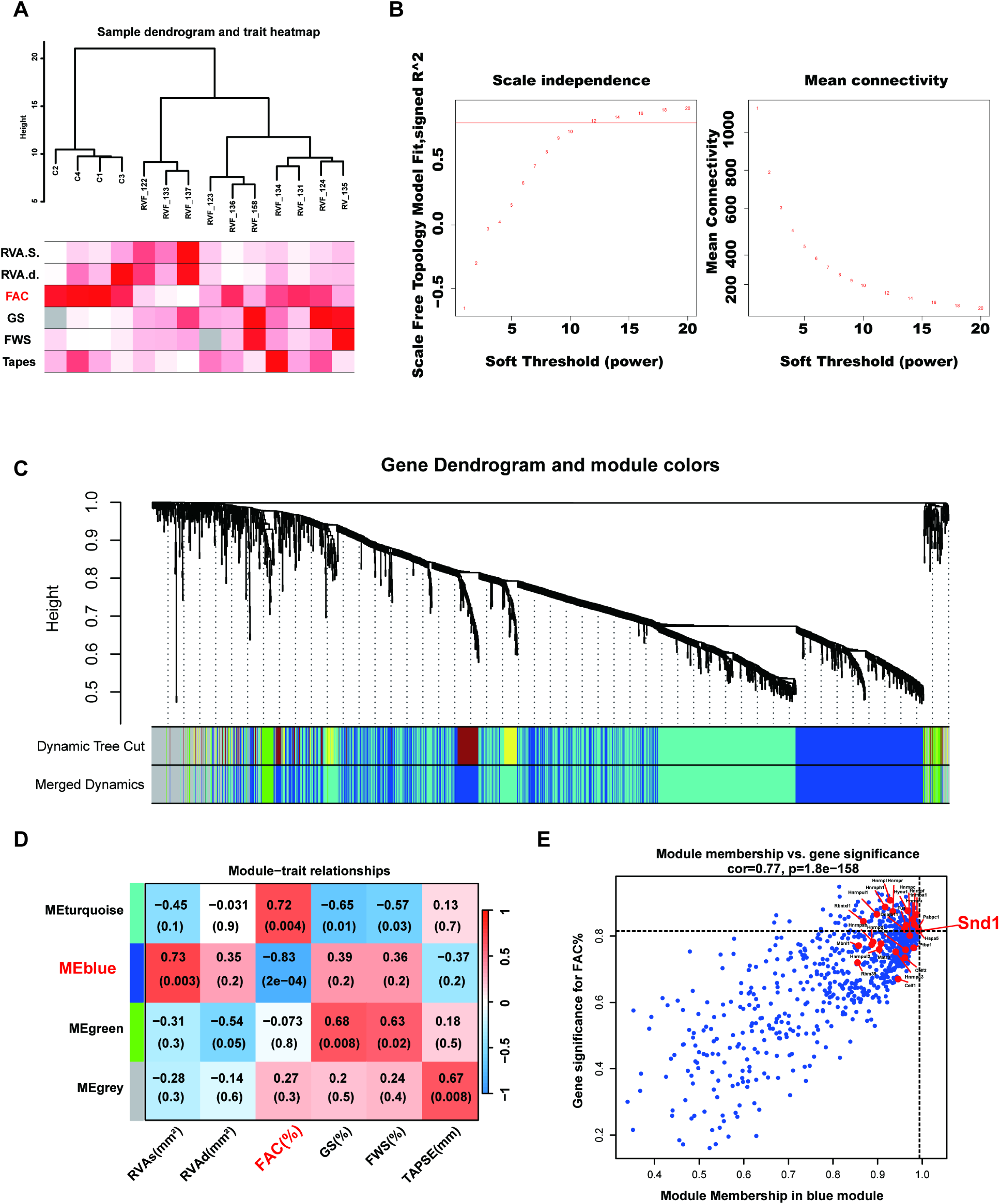
WGCNA reveals an association between high SND1 expression and the progression of RVF. **A.** Sample dendrogram and trait heatmap showing that fractional area change (FAC) differed most significantly between control and right ventricular failure (RVF) groups. **B.** A soft-thresholding power of 12 was selected based on scale-free topology and network connectivity to construct the weighted gene co-expression network. **C.** The gene clustering dendrogram based on topological overlap grouped genes into co-expression modules. Modules with an eigengene similarity greater than 0.75 were merged, resulting in four modules: blue, turquoise, green, and grey. **D.** Module-trait correlation heatmap showing the correlation between module eigengenes and different clinical data Each cell displays the correlation coefficient along with the corresponding P value. **E.** A scatter plot of gene significance (GS) versus module membership (MM) for genes in the MEblue module Previously reported fetal genes are highlighted in red. Among them, SND1 showed the highest module membership (MM = 0.993547) and strong gene significance (GS = 0.815517).

### SND1 is upregulated in failing cardiomyocytes

Correlation analysis between the relative protein levels of SND1 and FAC values in individual rats based on proteomic data revealed a significant negative correlation, indicating that elevated SND1 expression is associated with impaired RV function (Figure 3A). Through Western blot, we confirmed that SND1 was significantly upregulated in RVF myocardial tissue, accompanied by increased MYH7 and Caspase-3 and decreased SERCA2a and BCL-2 expression (Figures 3B–3G). Interestingly, no significant difference in SND1 mRNA levels was observed between the control and RVF groups (Figure 3H). Immunofluorescence further confirmed that SND1 upregulation occurred specifically in RV cardiomyocytes of RVF rats (Figure 3I). However, it is noteworthy that SND1 was not uniformly upregulated across all cardiomyocytes but showed abnormally high expression in a distinct subset (Figure S1A). No such phenomenon was observed in left ventricular tissues (Figure S1A). Further investigation into the role of SND1 in RVF progression and its underlying molecular mechanisms in cardiomyocyte impairment is required.

**Figure 3.**
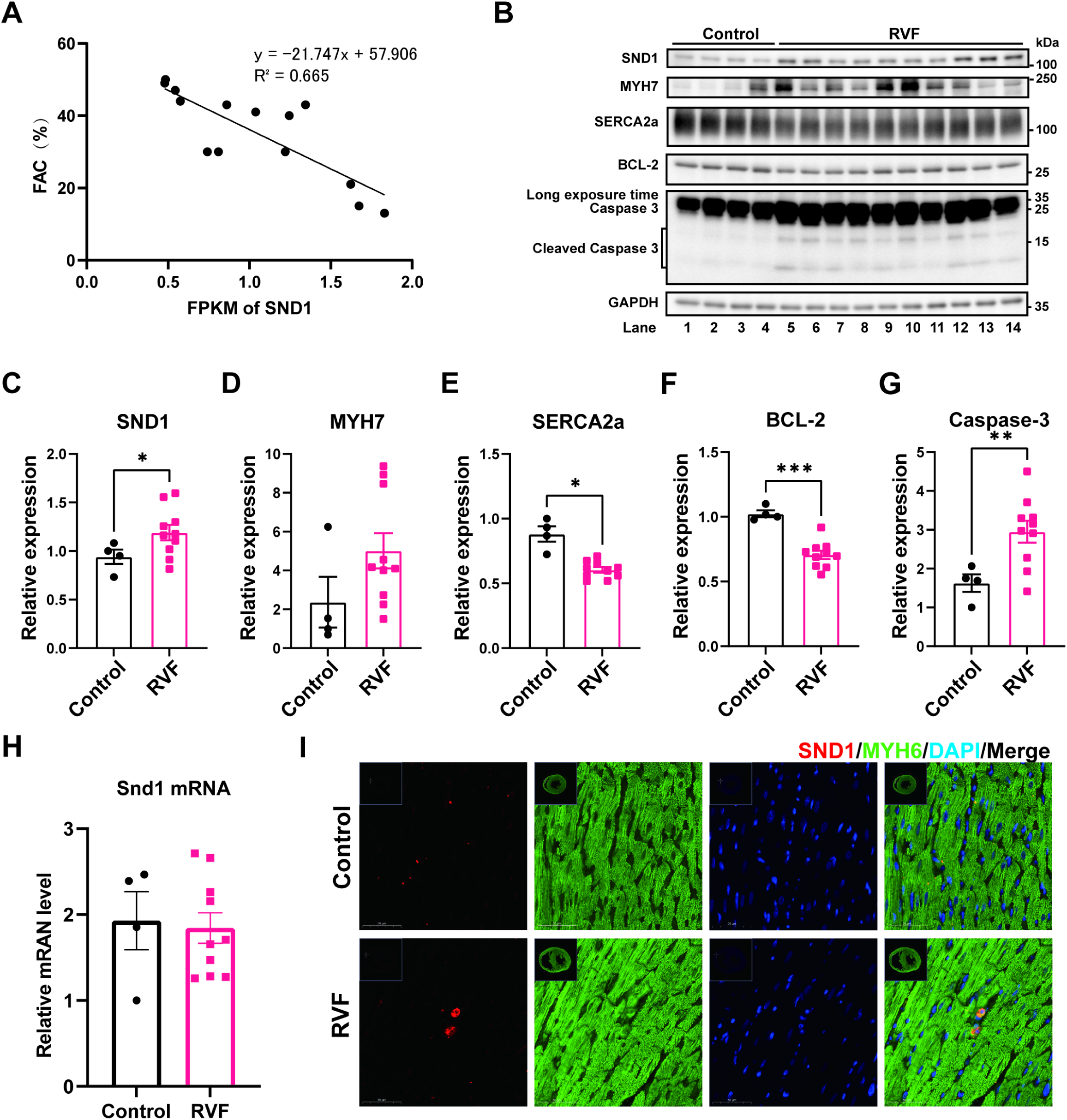
SND1 is upregulated in the right ventricular cardiomyocytes of RVF rats. **A.** A correlation analysis showing that the relative SND1 expression level detected by mass spectrometry is significantly associated with fractional area change (FAC), with an R² value of 0.665 **B–G.** SND1, MYH7, SERCA2a, BCL-2, and Caspase-3 protein levels in lysates from the right ventricles of control and RVF rats were analyzed by Western blotting with the indicated antibodies (n = 4–10). **H.** qPCR analysis showed no significant differences in SND1 mRNA levels in right ventricular tissue between control and right ventricular failure (RVF) groups. (n = 4–10). **I.** Immunofluorescence staining (red: SND1; green: MYH6; blue: DAPI) shows an upregulation of SND1 co-localized with MYH6 in cardiomyocytes. Scale bars: 50 μm All quantitative data are presented as the mean ± SEM. Statistical significance was determined using an unpaired Welch’s t-test. *P < 0.05, **P < 0.01, ***P < 0.001

### Neonatal and adult rat cardiomyocytes exhibit distinct SND1 protein levels and calcium-handling characteristics

Neonatal cardiomyocytes isolated from postnatal day 1 rats and adult rat cardiomyocytes from 8-week-old rats exhibit markedly distinct morphological and functional features. Neonatal cardiomyocytes display a characteristic spindle-shaped morphology with multiple protrusions, whereas adult cardiomyocytes appear more elongated and rod-shaped, with a relatively regular structure. Sarcomeric striations are faintly visible within adult cardiomyocytes, indicating an organized contractile architecture (Figure 4A). Previous studies reported that SND1 is a fetal gene whose expression is gradually silenced with heart maturity.^8^ Our experimental results confirmed this observation, showing that SND1 protein levels in the RV myocardium of neonatal rats were significantly higher than those in adult rats (Figures 4B and 4C). Immunofluorescence staining showed that SND1 is localized to the SR of neonatal cardiomyocytes, co-localizing with the SR protein SERCA2a (Figures 4D and 4E). Therefore, it may play a role in regulating SR function. The primary function of the SR in cardiomyocytes is calcium handling.^11^ An evaluation of calcium handling in isolated cardiomyocytes demonstrated that the baseline intracellular Ca²⁺ concentration in neonatal cardiomyocytes was significantly higher than in adult cardiomyocytes (Figures 4F and 4H). However, the velocity of Ca²⁺ release and reuptake during the calcium cycling process were lower in neonatal cardiomyocytes (Figures 4I and 4J). Comparison of normalized calcium transients also revealed that the amplitude of calcium transients was significantly reduced in neonatal cardiomyocytes compared with adult cardiomyocytes (Figures 4G and 4K). These findings raise the intriguing possibility that high SND1 expression and its localization to the SR may be associated with the impaired calcium handling capacity observed in neonatal cardiomyocytes compared to adult cardiomyocytes.

**Figure 4.**
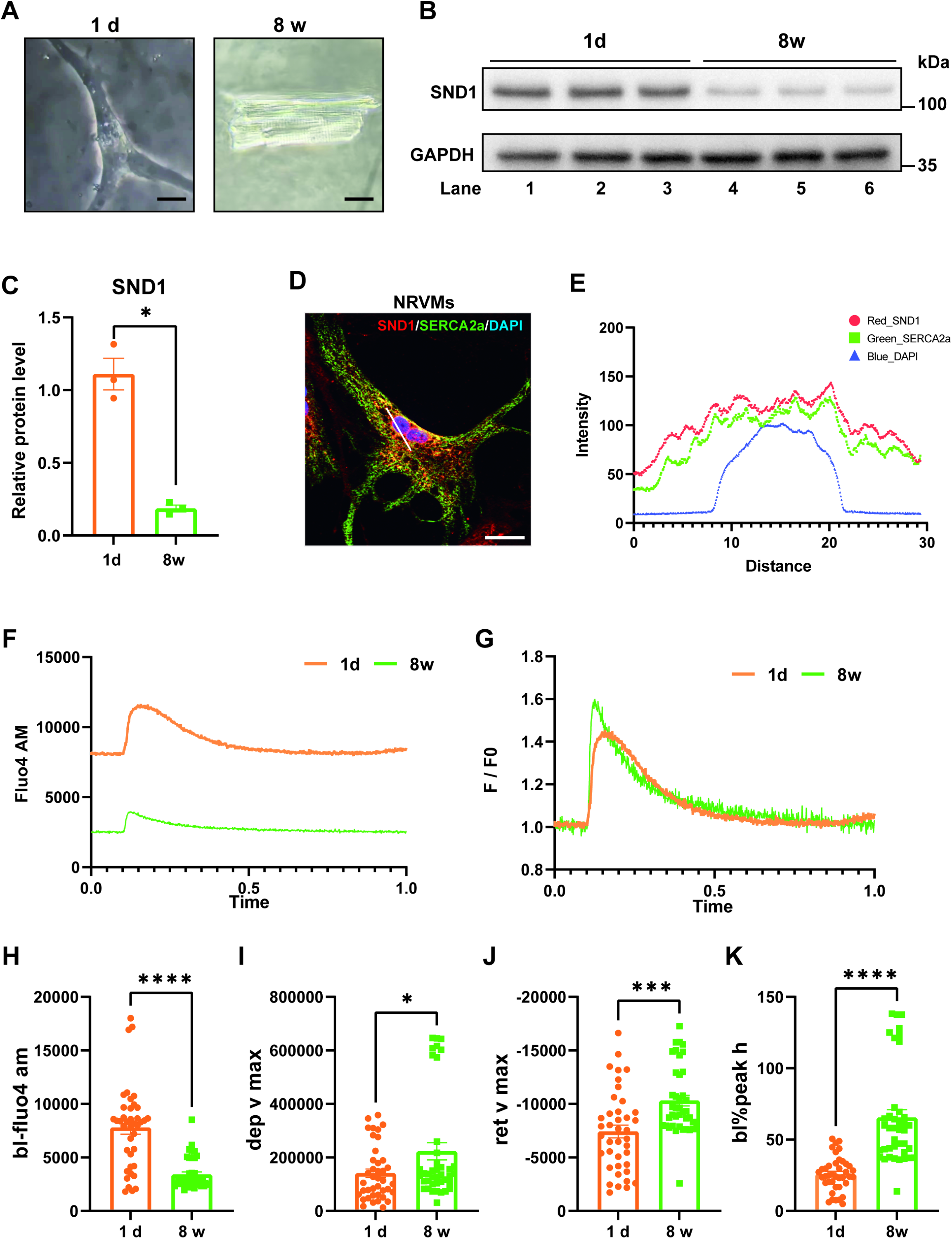
Differences in SND1 expression and calcium transient characteristics between neonatal and adult cardiomyocytes. **A.** Brightfield images showing morphological differences between neonatal myocytes and adult cardiomyocytes; scale bars: 20 μm **B–C.** SND1 protein levels in lysates from neonatal myocytes and adult cardiomyocytes were analyzed by Western blotting (n = 3). **D–E.** Immunofluorescence staining (red: SND1; green: SERCA2a; blue: DAPI) showed that SND1 is localized within the sarcoplasmic reticulum of NRVMs, showing clear colocalization with the SR marker SERCA2a. Scale bars: 20 μm **F.** Representative traces of raw calcium transient recordings **G.** Representative traces of normalized calcium transient recordings **H.** A Fluo4 AM baseline analysis of calcium transient was performed (n = 40 cells). **I.** Quantitative analysis of the maximal rate of Fluo4 AM during the calcium release phase of the transient (n = 40 cells) **J.** Quantitative analysis of the maximal rate of Fluo4 AM during the calcium recovery phase of the transient (n = 40 cells) **K.** Quantitative analysis of the percent change in calcium transient amplitude (n = 40 cells) All quantitative data are presented as the mean ± SEM. Statistical significance was determined using an unpaired Welch’s t-test. *P < 0.05, ***P < 0.001

### SND1 knockdown improves calcium handling and contractility in neonatal rat ventricular myocytes (NRVMs)

Given that SND1 is a cardiac fetal gene, it is naturally highly expressed in NRVMs. SND1 knockdown via siRNA in NRVMs allowed us to examine whether reducing SND1 expression could modulate SR-mediated calcium handling in cardiomyocytes (Figures 5A–5C). Fluo-4 AM staining revealed that SND1 knockdown significantly enhanced the velocity and amplitude of calcium transients in NRVMs without affecting baseline intracellular calcium levels (Figure 5D–5I). Furthermore, contraction tracing showed that the velocity of contraction and relaxation were significantly improved by SND1 knockdown in NRVMs (Figures 5J–5M). Together with our previous findings, these results suggest that SND1 may impair SR calcium handling, contributing to intracellular calcium overload and increased susceptibility to cardiomyocyte injury. Additionally, PI/Hoechst staining demonstrated a dose-dependent increase in NRVMs death following treatment with escalating concentrations of extracellular CaCl₂ (Figure 6A). Moreover, a Western blot analysis revealed a corresponding increase in Caspase-3 expression, indicating the activation of apoptotic pathways with calcium overload (Figure 6B). Knockdown of SND1 significantly attenuated CaCl₂-induced cell death (Figures 6C–6E) and suppressed the upregulation of Caspase-3 expression (Figures 6F–6H). These findings indicate that high SND1 levels may exacerbate calcium-induced cardiomyocyte injury, whereas SND1 knockdown could alleviate RV dysfunction by improving SR-mediated calcium cycling and reducing calcium-dependent cell death.

**Figure 5.**
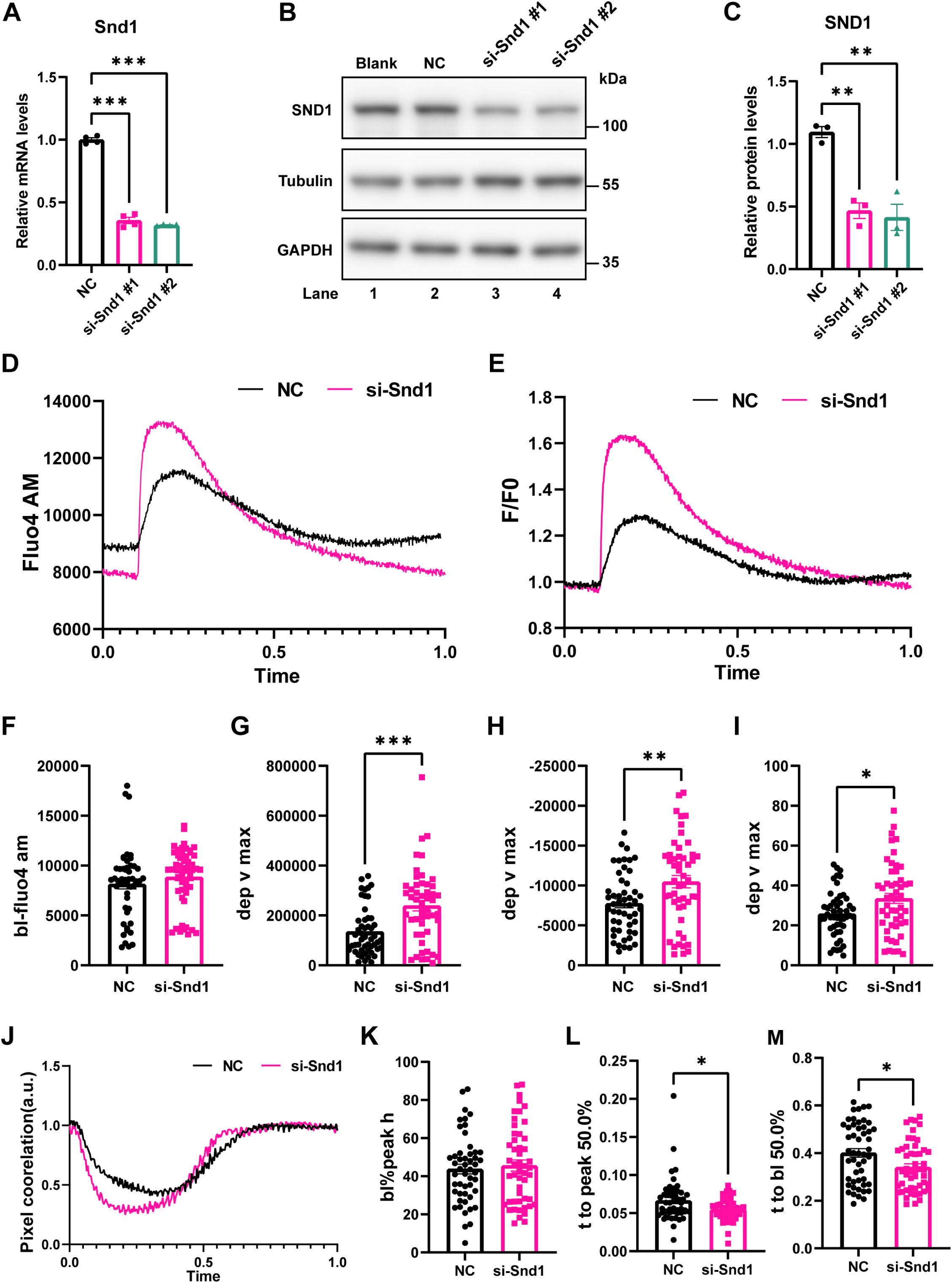
SND1 knockdown enhances contractility and calcium transients in neonatal rat ventricular myocytes (NRVMs). **A.** RT‒qPCR analyses of the mRNA levels of SND1 in NRVMs (n = 4) **B–C.** SND1 protein levels in lysates from NRVMs were analyzed by Western blotting (n = 3). **D.** Representative traces of raw calcium transient recordings **E.** Representative traces of normalized calcium transient recordings **F.** A Fluo4 AM baseline analysis of calcium transient was performed. **G.** Quantitative analysis of the maximal change in Fluo4 AM during the calcium release phase of the transient (n = 50 cells) **H.** Quantitative analysis of the maximal rate in Fluo4 AM during the calcium recovery phase of the transient (n = 50 cells) **I.** Quantitative analysis of the percent change in the calcium transient amplitude (n = 50 cells) **J.** Representative traces of the contraction curve of NRVMs **K.** Quantitative analysis of the percent change in the contraction amplitude (n = 50 cells) **L.** Quantitative analysis of the time to 50% of the contraction peak, a characterization of the speed of contraction (n = 50 cells) **M.** Quantitative analysis of the time to 50% of the baseline from the contraction peak, a characterization of the speed of cellular relaxation (n = 50 cells) All quantitative data are presented as the mean ± SEM. Statistical significance was determined using an unpaired Welch’s t-test. *P < 0.05, **P < 0.01, ***P < 0.001

**Figure 6.**
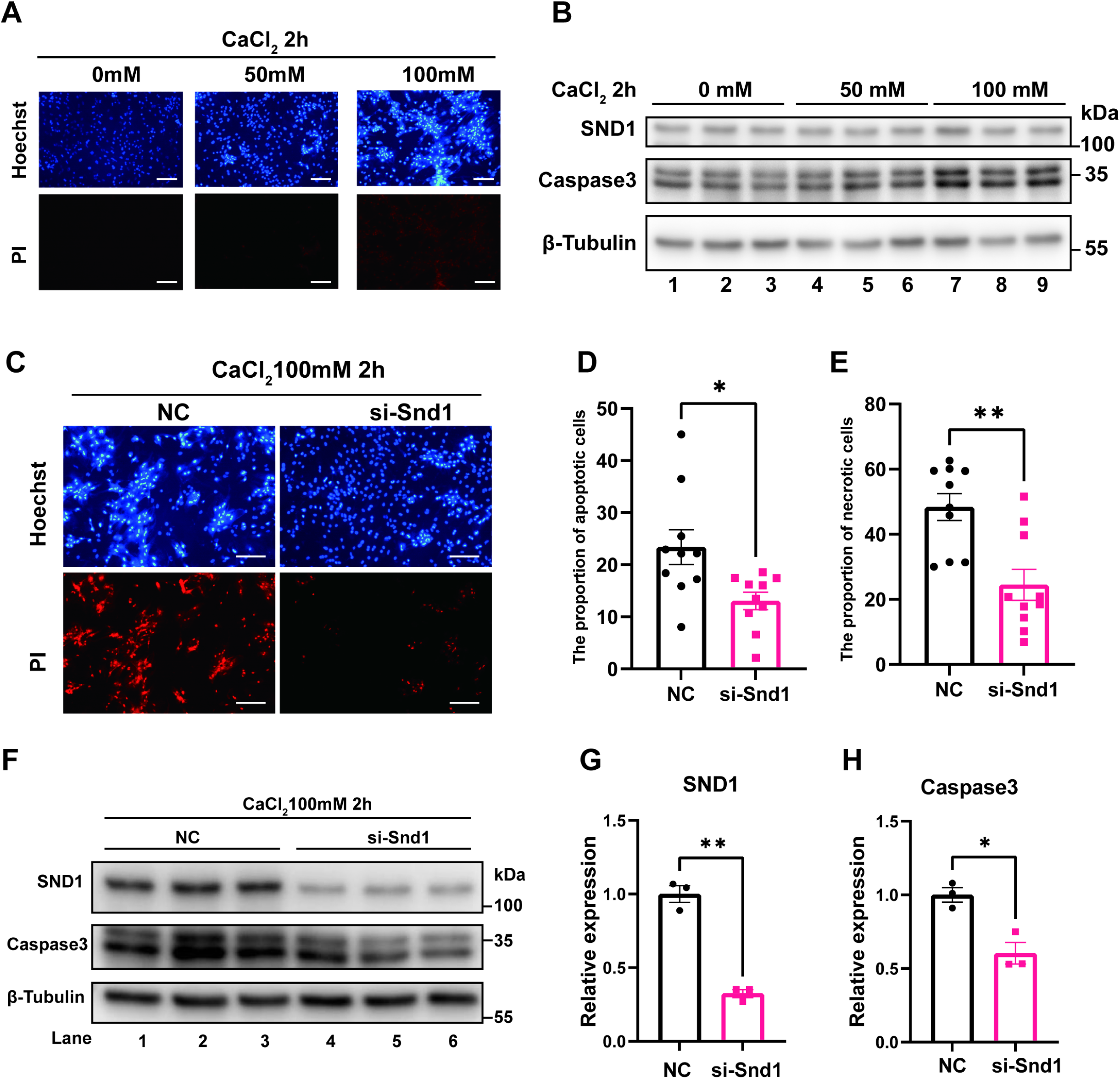
SND1 knockdown attenuates Ca²⁺ overload-induced cell death in neonatal rat ventricular myocytes (NRVMs). **A.** PI/Hoechst staining revealed a dose-dependent increase in NRVMs death with increasing extracellular Ca²⁺ concentrations in the culture medium. Scale bars: 100 μm **B.** SND1 and Caspase-3 protein levels in lysates from NRVMs were analyzed by Western blotting (n = 3). **C–E.** Quantitative analysis of PI/Hoechst staining; scale bars: 100 μm (n = 10) **F–H.** SND1 and Caspase-3 protein levels in lysates from NRVMs were analyzed by Western blotting (n = 3). All quantitative data are presented as the mean ± SEM. Statistical significance was determined using an unpaired Welch’s t-test. *P < 0.05, **P < 0.01

### SND1 interacts with SERCA2a and regulates its stability

To further elucidate the molecular mechanism by which SND1 regulates SR function, we performed co-immunoprecipitation coupled with mass spectrometry (Co-IP/MS) to identify SND1-interacting proteins in RVF myocardial tissue. Among the candidates, the key transporter regulating SR Ca^2+^ influx, SERCA2a, emerged as a potential target protein of SND1 (Figure 7A, Figure S2; Table S5). Preliminary verification of the mass spectrometry results was performed using protein samples from both NRVMs and RV tissues of RVF rats, through which we confirmed a physical interaction between SND1 and SERCA2a (Figures 7B and 7C). Furthermore, molecular docking analysis predicted the interaction interface and structural domains involved. The results suggested that SND1 may bind to SERCA2a via its SN3/SN4 domains. The interaction likely involves the amino acid region 644–677 of SERCA2a, with multiple potential hydrogen bonds stabilizing the interaction (Figures 7D and 7E).

**Figure 7.**
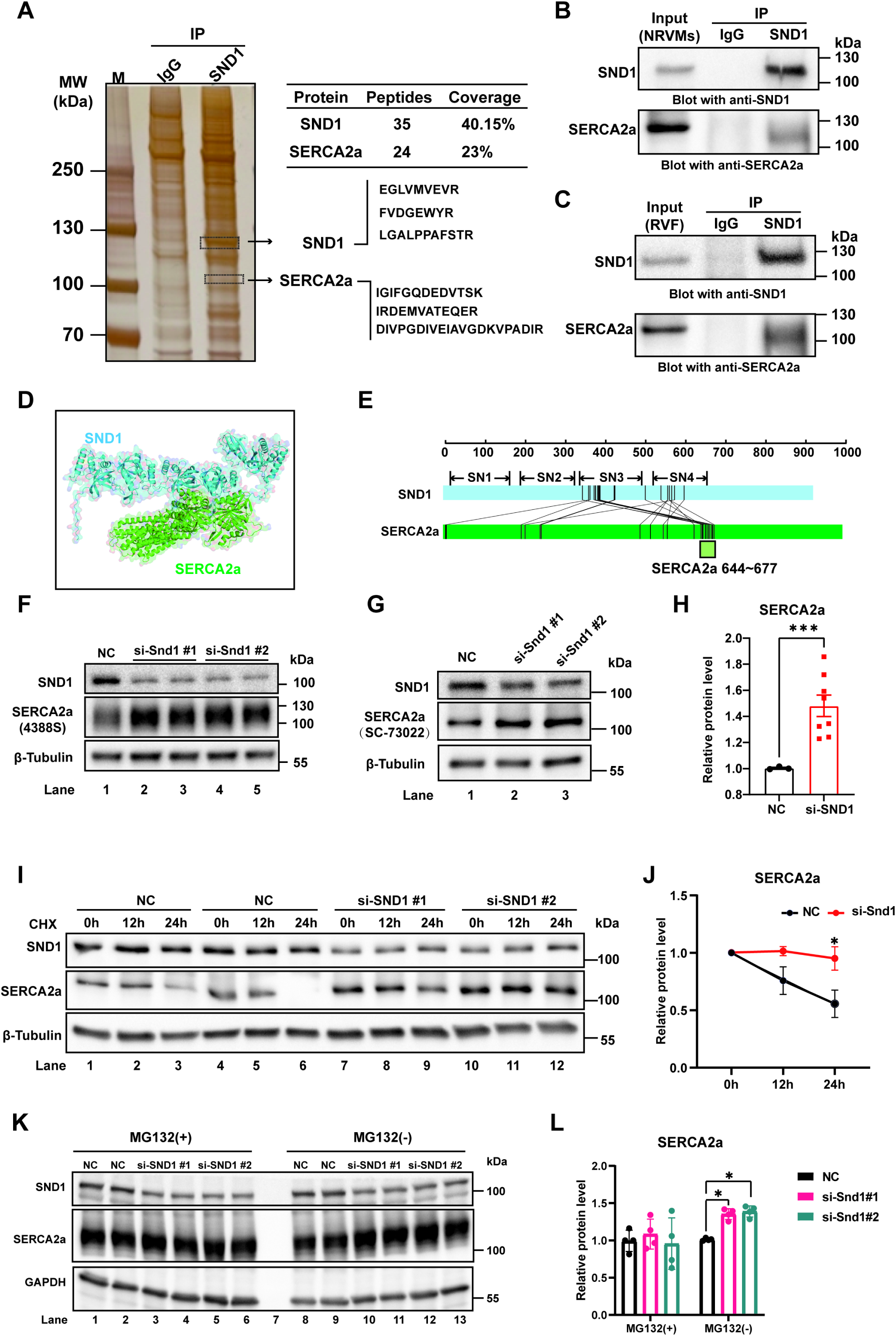
SND1 interacts with SERCA2a and promotes its degradation. **A.** Co-immunoprecipitation combined with liquid chromatography–mass spectrometry (LC-MS) identified an interaction between SND1 and SERCA2a in lysate extracts from the right ventricular (RV) myocardium of right ventricular failure (RVF) rats. **B.** Validation of the interaction between SND1 and SERCA2a in NRVMs **C.** Validation of the interaction between SND1 and SERCA2a in RV tissue samples of RVF rats **D.** Molecular docking analysis simulated the binding between SND1 (blue structure) and SERCA2a (green structure). **E.** Schematic illustration of the amino acid region through which SND1 primarily interacts with SERCA2a. **F–H.** SND1 knockdown and Western blot analysis of SERCA2a in NRVMs lysates were validated with two SERCA2a antibodies (n = 3–8). **I–J.** Western blot analysis of SND1 and SERCA2a in NRVMs lysates following cycloheximide (CHX) treatment in NC and si-SND1 groups (n = 3) **K–L.** Western blot analysis of SND1 and SERCA2a in NRVMs lysates following MG132 treatment in NC and si-SND1 groups (n = 4) All quantitative data are presented as the mean ± SEM. Statistical significance was assessed using an unpaired Welch’s t-test for comparisons between two groups, a two-way ANOVA for time-course experiments, and a one-way ANOVA for multiple-group comparisons. *P < 0.05, ***P < 0.001

The interaction between SND1 and SERCA2a suggests a potential regulatory role of SND1 in modulating SERCA2a. We observed a significant increase in SERCA2a protein levels following SND1 knockdown, which was not due to altered SERCA2a transcription (Figures 7F–7H and S3A). Thus, we hypothesized that SND1 might promote the intracellular degradation of SERCA2a. We then added cycloheximide (CHX) ed to prevent protein synthesis (Figure S3B). As expected, the rate of SERCA2a protein degradation significantly increased in SND1-knockdown cells (Figures 7I and 7J). Treatment with MG132 for 24 hours abolished the difference in SERCA2a protein levels between the SND1 knockdown and control groups, suggesting that SND1 may promote SERCA2a degradation via the proteasomal pathway (Figures 7K and 7L).

### SND1 CKO alleviates RV dysfunction under PAH

Our previous data indicated that upregulated SND1 in failing cardiomyocytes interacts with SERCA2a and promotes its degradation. Reduction in SERCA2a could impair SR calcium handling, thereby weakening myocardial contractility and increasing cell death due to calcium overload. Building on our findings, we set out to explicate the role of SND1 in PAH-induced RVF. We employed a tamoxifen-inducible Cre-LoxP system to achieve cardiomyocyte-specific SND1 deletion (SND1 CKO), enabling precise control over the timing of gene knockout. SND1^flox+/+^ and SND1^flox+/+^Myh6^cre+/-^ rats breed and develop normally and display no overt cardiovascular phenotype at baseline after tamoxifen administration (Figures S6–S8).

SND1 deletion alone did not affect baseline cardiac function. We, therefore, proceeded to investigate the impact of cardiomyocyte-specific SND1 knockout on the progression of RVF in PAH rats. First, littermate SND1^flox+/+^ and SND1^flox+/+^Myh6^cre+/-^ rats were genotyped on postnatal day 7 and grouped accordingly. Both groups were allowed free access to food and water and raised to 8 weeks of age, at which point PAH was induced via a single intraperitoneal injection of MCT. One week after the MCT injection, tamoxifen was administered intraperitoneally once daily for seven consecutive days to induce cardiomyocyte-specific SND1 knockout. Following tamoxifen administration, rats were maintained under normal feeding conditions for another two weeks to allow the rats to metabolize the residual tamoxifen. All remaining rats, apart from those used in the survival analysis, underwent transthoracic echocardiography and harvesting of RV tissues on day 28 post-MCT injection (Figure 8A).

**Figure 8.**
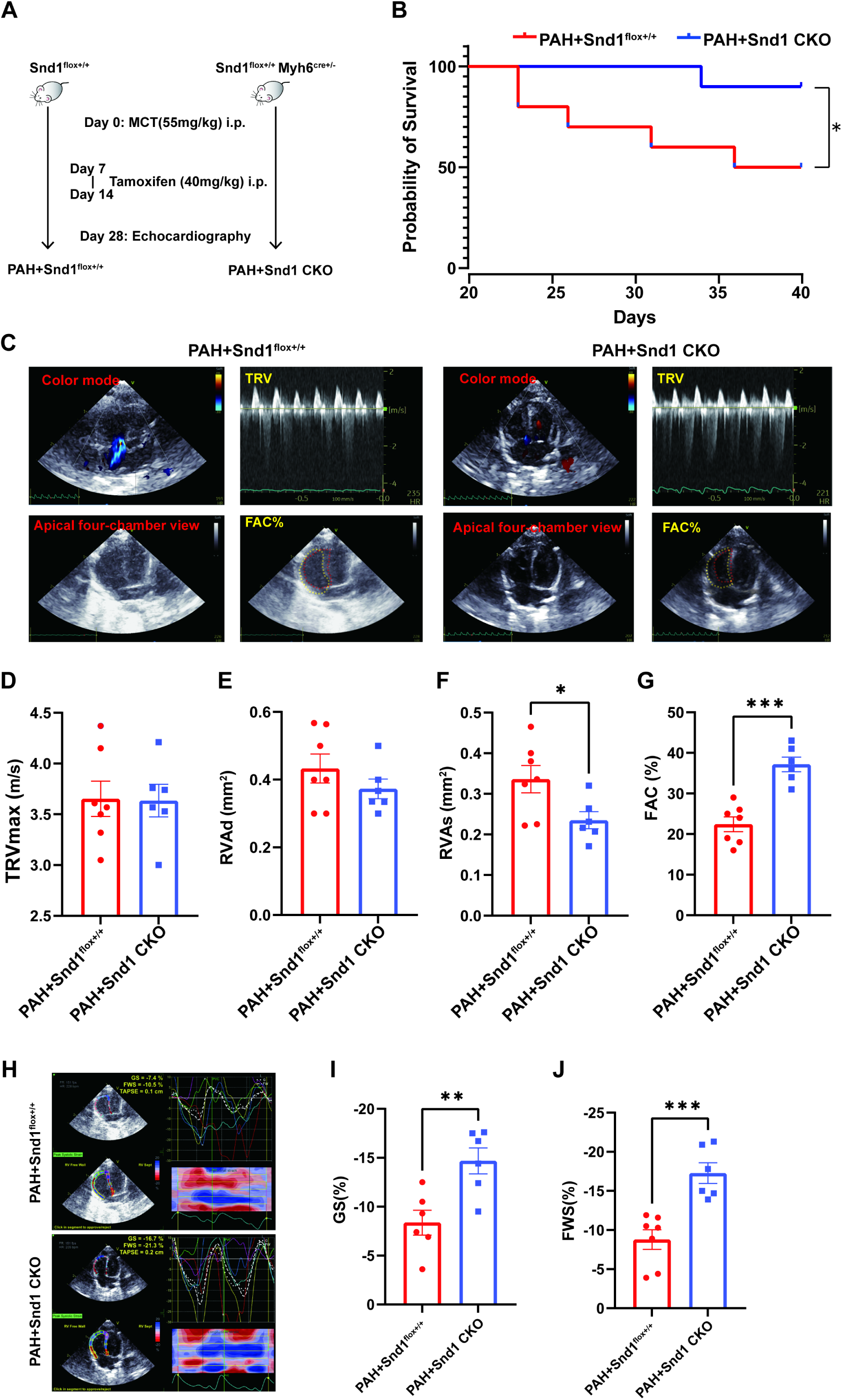
Cardiomyocyte-specific deletion of SND1 improves right ventricular function and prolongs survival in pulmonary arterial hypertension (PAH) rats. **A.** Schematic diagram of the experimental protocol Age- and weight-matched rats received a single intraperitoneal injection of monocrotaline (MCT) to induce PAH. Tamoxifen was administered intraperitoneally from days 7 to 14 to induce SND1 deletion. All the rats underwent transthoracic echocardiography and tissue collection on day 28 (PAH + Snd1^flox+/+^ vs PAH + Snd1 CKO), except for those used for the survival analysis. **B.** Kaplan–Meier survival analysis showed that SND1 CKO significantly improved survival in PAH rats. The Log-rank test was used to assess statistical significance (n = 10, *P < 0.05). **C.** Representative images of transthoracic echocardiography performed to evaluate tricuspid regurgitation velocity (TRV) and right ventricular fractional area change (FAC) **D.** Quantitative analysis of the TR peak velocity, a characterization of the right ventricular pressure load (n = 6–7) **E.** Quantitative analysis of the end-diastolic area (RVAd) (n = 6–7) **F.** Quantitative analysis of the end-systolic area (RVAs) (n = 6–7) **G.** Quantitative analysis of FAC, a characterization of right ventricular function (n = 6) **H.** Representative images of right ventricular strain using speckle-tracking echocardiography I. Quantitative analysis of global strain (GS) (n = 6) **J.** Quantitative analysis of free wall strain (FWS) (n = 6–7) All quantitative data are presented as the mean ± SEM. Statistical comparisons between groups were performed using unpaired Welch’s t-tests. *P < 0.05, **P < 0.01, ***P < 0.001

Kaplan–Meier survival analysis demonstrated that SND1 CKO significantly improved survival in PAH rats (Figure 8B). On day 28 after MCT administration, both groups of rats underwent transthoracic echocardiography (Figure 8C). Measurement of the TR peak velocity (TRVmax) revealed no significant difference between PAH + SND1^flox+/+^ and PAH + SND1 CKO rats, suggesting that both groups had a comparable degree of RV afterload (Figure 8D). Under equivalent pressure load, PAH + SND1 CKO rats had significantly reduced RVAs and a correspondingly increased FAC compared to control PAH rats (Figure 8E-8G). Furthermore, speckle-tracking echocardiography revealed that RV myocardial strain was significantly improved in PAH + SND1 CKO rats compared to PAH + SND1^flox+/+^ rats (Figures 8H–8J). These results indicate that cardiomyocyte-specific SND1 deletion improves RV systolic function in PAH rats.

PV loop analysis confirmed that the RV pressure load was comparable between the two groups. However, ventricular dilation induced by elevated pressure overload was significantly attenuated in PAH + SND1 CKO rats. End-systolic and end-diastolic RV volumes were markedly lower in the PAH + SND1 CKO group than in the PAH + SND1^flox+/+^ group. In the PV loop experiment, transient occlusion of the inferior vena cava was used to reduce venous return and thereby decrease cardiac preload; this allowed us to assess the linear relationship between RV pressure and volume. The slope of this relationship (end-systolic elastance [Ees]) reflects myocardial contractility and is positively correlated with ventricular systolic function. A comparison of RV Ees between PAH + SND1^flox+/+^ and PAH + SND1 CKO rats revealed that RV contractility was significantly enhanced in the SND1 CKO group. These findings demonstrate that SND1 CKO enhances the ability of the right ventricle to preserve its structural integrity under high-pressure load, primarily by improving myocardial contractility.

Histological analysis of cardiac tissue confirmed that SND1 CKO markedly attenuated adverse RV remodeling and cardiomyocyte apoptosis in PAH rats. H&E staining revealed that in PAH + SND1^flox+/+^ rats, the RV free wall had disorganized myocardial fiber alignment, hyperchromatic and distorted nuclei, and compensatory hypertrophy with structural distortion in the lower interventricular septum. These pathological features were significantly improved in PAH + SND1 CKO rats (Figure 9G). In addition, TUNEL staining showed a significantly lower proportion of TUNEL-positive nuclei in the RV tissues of PAH + SND1 CKO rats than in those of PAH + SND1^flox+/+^ rats, indicating reduced cardiomyocyte apoptosis (Figure 9H). A Western blot analysis confirmed that SND1 CKO reduced apoptosis in the myocardial tissue of PAH rats, as evidenced by decreased Caspase-3 protein levels. Moreover, SND1 CKO also reversed the downregulation of SERCA2a, consistent with the conclusions drawn from the in vitro experiments (Figures 9I–9K). These findings highlight the therapeutic potential of targeting SND1 to preserve myocardial structure and function in RVF.

**Figure 9.**
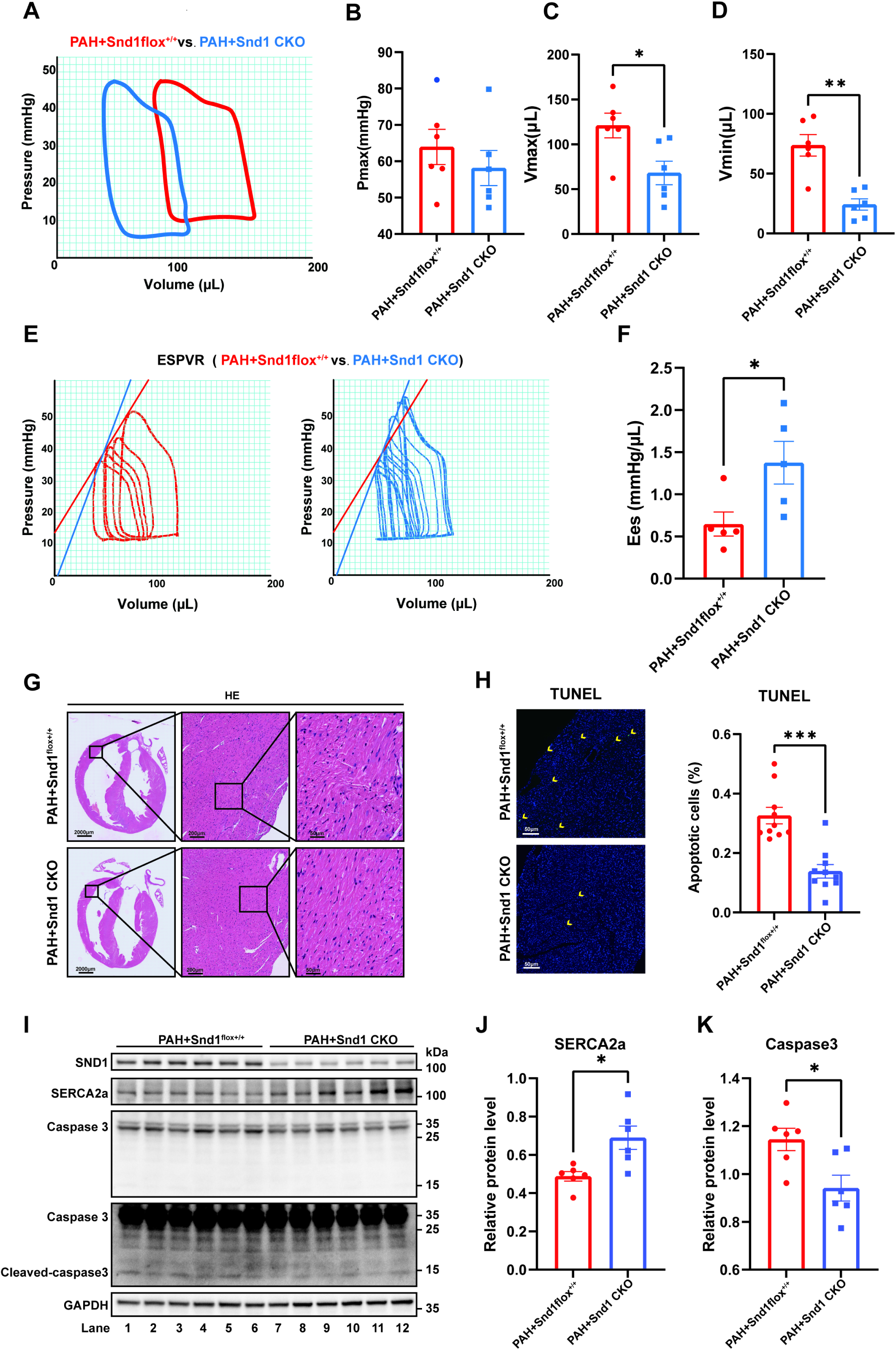
Cardiomyocyte-specific deletion of SND1 enhances myocardial contractility, attenuates adverse remodeling, upregulates SERCA2a expression, and reduces apoptosis in PAH rats. **A.** Right heart catheterization was performed to record pressure-volume (PV) loops of the right ventricle. **B.** Quantitative analysis of pressure measurements in the right ventricle (n = 6) **C–D.** Quantitative analysis of volume measurements in the right ventricle (n = 6) **E.** PV loops were used to evaluate myocardial contractility by calculating the end-systolic pressure-volume relationship (ESPVR). The slope of this relationship, known as the end-systolic elastance (Ees), reflects contractile function. **F.** Quantitative analysis of the Ees of the right ventricle (n = 5) **G.** Paraffin-embedded sections of heart tissue were subjected to HE staining. The boxed area was further enlarged. Scale bars: 2000 μm, 200 μm, and 50 μm (from left to right) **H.** Quantitative analysis of TUNEL staining by the proportion of apoptotic nuclei in right ventricular tissue. Scale bars: 50 μm (n = 10) **I–K.** SND1, SERCA2a, and Caspase-3 protein levels in lysates from right ventricular tissue were analyzed by Western blotting (n = 5). All quantitative data are presented as the mean ± SEM. Statistical comparisons were performed using unpaired Welch’s t-tests. *P < 0.05

## Discussion

To verify the fetal gene characteristics of SND1, we compared SND1 protein levels in RV tissue from neonatal and healthy adult rats. Consistent with the findings reported by Professor Jie Yang’s group,^8^ SND1 expression was significantly downregulated in the right ventricle of 8-week-old rats as the heart matured. These results indicate that in contrast to healthy adult cardiomyocytes, neonatal and failing cardiomyocytes exhibit high levels of SND1 expression.

Therefore, NRVMs provide a theoretically and practically feasible cell model with naturally high SND1 expression to investigate the functional role and molecular mechanisms of SND1 in cardiomyocytes. Immunofluorescence co-staining revealed that SND1 is predominantly localized to the SR in NRVMs, showing strong colocalization with SR marker proteins. This observation provides important mechanistic clues, suggesting SND1 may contribute to cardiomyocyte injury in heart failure by regulating SR function. The SR is a critical organelle for maintaining calcium homeostasis in cardiomyocytes, essential for the rhythmic contraction and relaxation of the heart.^11^ During each cardiac cycle, depolarization of the cardiomyocyte membrane allows extracellular Ca²⁺ influx into the cytosol through L-type calcium channels located on the transverse tubules. This triggers the opening of RyR2 receptors on the SR membrane, resulting in a rapid release of stored Ca²⁺ into the cytosol. The sharp rise in cytosolic Ca²⁺ concentration allows calcium to bind troponin C, initiating sarcomere contraction. Subsequently, most cytosolic Ca²⁺ is rapidly pumped back into the SR, while the sodium-calcium exchanger (NCX) on the plasma membrane extrudes a smaller portion. This reuptake restores a steep Ca²⁺ gradient between the SR and cytosol, critical for preparing the cell for the next contraction. Disruption of SR function leads to cytosolic calcium overload, which blunts calcium transients and reduces myocardial contractility.^12^ Compared with adult cardiomyocytes, NRVMs naturally express higher levels of SND1. We observed that both the amplitude and velocity of calcium transients were significantly higher in adult cardiomyocytes than in NRVMs. Interestingly, a similar difference in calcium handling was observed in NRVMs before and after SND1 knockdown, supporting our hypothesis that high levels of SND1 may suppress SR function. Thus, we propose that SND1 inhibits SR-mediated calcium handling and that its knockdown enhances SR function. This increases the calcium transient amplitude and velocity, ultimately improving cardiomyocyte contractility.

In addition, cytosolic calcium overload acts as a secondary messenger to activate various calcium-dependent signaling pathways, contributing to pathological cardiomyocyte hypertrophy, mitochondrial dysfunction,^13^ endoplasmic reticulum (ER) stress, and apoptosis.^14,15^ Calcineurin, a calcium-dependent phosphatase, is significantly activated in response to elevated intracellular Ca²⁺ levels. One of its downstream effects is the dephosphorylation of Drp1 at serine 637 (Ser637), which promotes the recruitment of Drp1 to the mitochondria and subsequently triggers mitochondrial fission.^16^ In parallel, calcineurin also dephosphorylates the transcription factor NFAT (nuclear factor of activated T-cells), facilitating its translocation from the cytosol to the nucleus, where it induces the expression of pro-hypertrophic genes.^17^ Moreover, calmodulin (CaM) binds Ca²⁺ and activates Ca²⁺/CaM-dependent protein kinase II (CaMKII), whose sustained activation is a key driver of cardiac pathologies, including maladaptive remodeling, arrhythmogenesis, and cardiomyocyte death.^18^ To experimentally model calcium overload, we treated NRVMs with a culture medium containing high concentrations of CaCl₂. The excess extracellular Ca²⁺ entered the cardiomyocytes through calcium channels, leading to intracellular calcium overload and apoptosis. Notably, SND1 knockdown significantly attenuated CaCl₂-induced apoptosis in NRVMs treated with 100 μM CaCl₂, highlighting the therapeutic potential of targeting SND1 under calcium overload conditions. These findings further support our hypothesis that elevated SND1 impairs SR function, exacerbating calcium dysregulation and contributing to cardiomyocyte injury.

From a structural biology perspective, SND1 is a multi-domain protein that contains several staphylococcal nuclease (SN)-like domains and a Tudor domain.^19,20^ The SN domains belong to the oligonucleotide/oligosaccharide-binding (OB)-fold superfamily, which is evolutionarily conserved. Each SN domain consists of a characteristic OB-fold composed of a five-stranded β-barrel flanked by three α-helices on each side.^21–23^ The Tudor domain, a conserved structural motif of approximately 60 amino acids, comprises five antiparallel β-strands that form a sharply curved β-sheet structure, giving rise to a barrel-like fold.^24^ Structurally, the four N-terminal SN domains of SND1 form a rod-like shape, while the C-terminal Tudor domain, partially fused with other SN domains, adopts a hook-like conformation. This multi-domain architecture enables SND1 to interact with nucleic acids, individual proteins, and protein complexes in diverse ways, allowing it to regulate multiple cellular signaling pathways and influence disease progression.^20,25^ For example, SND1 was originally identified as a transcriptional coactivator capable of interacting with transcription factors such as STAT6 and c-Myb. In addition, SND1 is involved in post-transcriptional processes, including spliceosome assembly, microRNA (miRNA) degradation, and RNA stability regulation.^20,26–28^ Functionally, the SN3 domain of SND1 interacts with HLA-A, promoting immune evasion in tumor cells,^29^ while the SN1/SN2 domains bind metadherin (MTDH) and enhance its oncogenic activity.^30^ Therapeutic targeting of these interactions has been shown to alleviate disease progression in relevant models.^26,31^ These findings suggest that SND1 is not merely a passive modulator of gene expression but may be a key functional driver under pathological conditions, with protein-protein interactions serving as a critical mechanism for its biological activity. In our study, we performed CoIP/MS to identify the proteins interacting with SND1 in the RV myocardium of RVF rats. Among the candidates, we identified SERCA2a, an SR Ca²⁺-ATPase essential for calcium handling, as a potential SND1-interacting partner. We conducted repeated CoIP–Western blot experiments in both RVF rat RV tissue and NRVMs to validate this interaction. We consistently detected a physical interaction between SND1 and SERCA2a in both systems. Based on these findings, SND1 may impair SR function in cardiomyocytes by interacting with and modulating SERCA2a, contributing to calcium handling dysfunction in the failing right ventricle.

In mammals, the sarcoplasmic/endoplasmic reticulum Ca²⁺-ATPase (SERCA) family consists of multiple protein isoforms encoded by SERCA1, SERCA2, and SERCA3. Among these, SERCA2a is the predominant isoform expressed in cardiomyocytes and type I skeletal muscle fibers.^32^ As a key regulator of calcium cycling in cardiomyocytes, SERCA2a is responsible for the reuptake of cytosolic Ca²⁺ into the SR (SR), thereby maintaining intracellular calcium homeostasis. Following Ca²⁺-induced sarcomere contraction, SERCA2a pumps Ca²⁺ back into the SR in an ATP-dependent manner, enabling cardiac relaxation.^10^ The SERCA2a expression level and functional activity are crucial for maintaining calcium balance in cardiomyocytes and are directly linked to cardiac contractility. A substantial body of research has identified SERCA2a as a pivotal determinant of cardiomyocyte fate and a central player in heart failure progression.^10^ Reduced expression or activity of SERCA2a has been associated with both systolic and diastolic dysfunction in human heart failure.^33,34^ Various post-translational modifications have been shown to modulate SERCA2a activity and contribute to heart failure progression. Changes in SERCA2a protein abundance have been directly linked to deteriorating cardiac function. Complete genetic ablation of SERCA2a is lethal to embryos, while SERCA2 heterozygous (+/–) knockout mice exhibit reduced SR Ca²⁺ uptake and impaired contractile function^35^. Moreover, conditional cardiomyocyte-specific deletion of SERCA2a results in decreased cytosolic Ca²⁺ clearance, reduced SR Ca²⁺ content, diminished Ca²⁺ transients, and severe heart failure within weeks after gene deletion.^36–38^ Notably, restoring deubiquitinase activity alleviates SERCA2a degradation and the associated cardiac dysfunction.^40^ The present study identified SND1 as a novel regulator of SERCA2a protein abundance in cardiomyocytes. Our data suggest that SND1 may promote proteasomal degradation of SERCA2a through direct protein-protein interaction, representing a previously unrecognized mechanism contributing to calcium handling imbalance and SERCA2a loss in heart failure. Based on the findings of this study, it is conceivable that targeting the protein-protein interaction between SND1 and SERCA2a may be a promising therapeutic strategy. The development of small-molecule inhibitors or peptide-based disruptors that interfere with this interaction could potentially mitigate proteasomal degradation of SERCA2a, thereby preserving calcium homeostasis and improving myocardial contractility in heart failure. This approach holds potential translational value for the treatment of RV dysfunction in PAH and other forms of heart failure.

Several important considerations were made in the design of our animal experiments. First, although there is no published evidence to date because SND1 is a newly identified cardiac fetal gene, global knockout may lead to impaired cardiac development or even embryonic death. Therefore, we employed a cardiomyocyte-specific, conditional knockout strategy using the flox-Cre system, allowing for spatial and temporal control of SND1 deletion. Second, it is well-established that estrogen confers cardioprotective effects through multiple mechanisms. These include enhancement of cardiomyocyte fatty acid metabolism and mitochondrial function, reduction of oxidative stress, inhibition of collagen deposition, attenuation of hypertrophic responses, and anti-apoptotic effects.^40–43^ Including female rats in the experimental groups could result in substantial intra-group variability due to estrogen-related protection, thereby increasing the risk of false-negative conclusions. Additionally, we used Myh6 CreERT2 transgenic rats as the Cre driver line. Although the CreERT2 fusion protein is theoretically inactive in the cytoplasm without tamoxifen induction, our preliminary findings showed that female SND1^flox+/+^ Myh6^Cre+/-^ rats weighed approximately half the body weights of their male littermates, a difference far beyond the expected physiological sex-based range. We suspect that endogenous estrogen in female rats may have partially activated the CreERT2 system, leading to premature SND1 recombination and affecting normal development. This represents a potential confounding factor. Therefore, we primarily selected male rats for the in vivo experiments to ensure consistency and minimize variability. Finally, in our in vitro experiments, SERCA2a protein levels, cardiomyocyte contractility, and cell viability emerged as the three primary outcome measures linked to SND1-related calcium handling dysfunction. Consequently, in the in vivo validation phase, we focused on evaluating the impact of SND1 conditional knockout on these specific endpoints in the RV myocardium of PAH rats. Nevertheless, we acknowledge the possibility that SND1 deletion may improve cardiac outcomes through additional mechanisms that are not directly related to calcium homeostasis.

This study has certain limitations. Although we report for the first time an interaction between SND1 and SERCA2a and provide preliminary evidence suggesting that SND1 may promote the degradation of SERCA2a, thereby impairing SR calcium reuptake and subsequently disrupting calcium homeostasis and contractile function in cardiomyocytes, several aspects remain to be clarified. Specifically, our analysis of the interaction domains between SND1 and SERCA2a was limited to in silico predictions, and no structural validation or in-depth biochemical mapping was performed. Moreover, the causal relationship between SND1-SERCA2a interaction, SERCA2a degradation, and calcium handling dysfunction is inferred from our current experimental findings, but direct mechanistic evidence is still lacking.

In summary, this study elucidates a previously unrecognized molecular mechanism underlying calcium homeostasis imbalance in RVF. Our findings suggest that cardiac fetal gene reprogramming contributes to disrupted calcium handling in failing cardiomyocytes, exacerbating RV dysfunction and apoptosis in a rat model of PAH. Specifically, the fetal gene product SND1 is upregulated in RVF cardiomyocytes, where it interacts with the SR Ca²⁺-ATPase SERCA2a and promotes its degradation via the proteasomal pathway. This interaction impairs SR-mediated reuptake of cytosolic calcium, leading to reduced contractility, weakened calcium transients, and increased susceptibility to calcium overload-induced cell death. Knockdown or genetic deletion of SND1 mitigates SERCA2a degradation, enhances cardiomyocyte contractile performance and calcium handling, and improves cellular tolerance to calcium stress. These findings suggest that SND1 may be a potential therapeutic target for alleviating RV dysfunction by restoring calcium homeostasis in the failing heart.

## Acknowledgments

None.

## Funding

This work was supported by the National Natural Science Foundation of China (Grant No. 82170291, 32270722), National Natural Science Foundation Regional Innovation and Development Joint Fund (Grant No. U24A20230), and Natural Science key Foundation of Hunan Province (Grant No. 2024JJ3051).

## Author contributions

Conceptualization: Lizhe Guo, E Wang, Jian Qiu; Methodology: Lizhe Guo, Xingyang Liu, Lu Wang, Dandan Wang, Xu Cheng; Formal analysis and investigation: Lizhe Guo, Xiaocheng Zhu; Chunyan Ye, Na Chen, Longyan Li, Yanfeng Wang; Writing - original draft preparation: Lizhe Guo, E Wang; Writing - review and editing: Yinqi Li, Gang Qin, Jian Qiu; Supervision: E Wang, Jian Qiu.

## Availability of data and materials

The data presented in this study are available on request from the corresponding authors.

## Ethical approval

All animal experiments were carried out in accordance with the recommendations of national and international animal care and ethical guidelines and were approved by the Ethics Committee for Animal Research of Xiangya Hospital of Central South University (permit code: 202309005). The manuscript does not contain clinical studies or patient data.

## Consent for publication

Yes.

## Conflict of interest

None.

## Non-standard abbreviations and acronyms

ER: Endoplasmic reticulum
ESPVR: End-systolic pressure-volume relationship
FAC: Fractional area change
FWS: Free wall strain
GS: Global strain
MM: Module membership
NFAT: Nuclear factor of activated T-cells
NRVMs: Neonatal rat ventricular myocytes
PAH: Pulmonary arterial hypertension
PV: Pressure-volume
RV: Right ventricular
RVA: Right ventricular end-systolic area
RVF: Right ventricular failure
SR: Sarcoplasmic reticulum
TOP: Terminal oligo-pyrimidine
TR: Tricuspid regurgitation
WGCNA: Weighted gene co-expression network analysis
H&E: Hematoxylin and Eosin
TUNEL: Terminal deoxynucleotidyl transferase dUTP Nick-End Labeling

## Notes

### Competing Interest Statement

The authors have declared no competing interest.

